# Phage steering of antibiotic-resistance evolution in the bacterial pathogen *Pseudomonas aeruginosa*

**DOI:** 10.1101/868661

**Authors:** James Gurney, Léa Pradier, Joanne S. Griffin, Claire Gougat-Barbera, Benjamin K. Chan, Paul E. Turner, Oliver Kaltz, Michael E. Hochberg

## Abstract

Antimicrobial resistance is a growing global concern and has spurred increasing efforts to find alternative therapeutics. Bacteriophage therapy has seen near constant use in eastern Europe since its discovery over a century ago. One promising approach is to use phages that not only reduce bacterial pathogen loads, but also select for phage resistance mechanisms that trade-off with antibiotic resistance – so called ‘phage steering’. Recent work has shown that phage OMKO1 can interact with efflux pumps and in so doing select for both phage resistance and antibiotic sensitivity. We tested the robustness of this approach to three different antibiotics *in vitro* and one *in vivo.* We show that *in vitro* OMKO1 can reduce antibiotic resistance either in the absence or the presence of antibiotics. Our *in vivo* experiment showed that phage increased the survival times of wax moth larvae and increased bacterial sensitivity to erythromycin, both in the absence and presence of the antibiotic. We discuss the implications of our findings for future research on this promising therapeutic approach using OMKO1.

## Introduction

Increasing levels of antimicrobial resistance have led to the exploration of alternative treatments and strategies (1, 2). Lytic bacteriophage (phage) has demonstrated therapeutic potential, particularly for recalcitrant bacterial infections (3). Phage therapy works in a similar way to conventional chemotherapies – bacterial densities are reduced to the extent that the immune system can clear the infection (4-7). However, as with conventional therapies, treatment may fail if phages are not able to sufficiently reduce bacterial density, often attributable to bacteria evolving phage resistance during therapy. Many strategies have been proposed that go beyond simple phage monotherapy with the aim of preventing bacterial resistance. These capitalize on ecological and evolutionary events in the disease ecosystem and include multi-phage cocktails, evolutionarily trained ‘sur-mesure’ phage, engineered phage, or combining phages with antibiotics (1).

One particularly promising approach is to anticipate resistance evolution to phages and use this as part of a therapeutic strategy, recently termed ‘phage steering’ (8). This involves exploiting an evolutionary trade-off between resistance to the phage and either sensitivity to other antimicrobials (a ‘double bind’ (9)) or reduced bacterial virulence (2, 10). Several phage resistance mechanisms have been identified that generate such trade-offs, one of the most promising being the reduction in efflux pump efficiency associated with phage resistance (3, 11). This form of phage steering has the potential to prevent the emergence of antibiotic resistance or even revert a resistant bacterial infection to antibiotic sensitivity (8).

Several candidate phages have been identified for phage steering and are currently being used as compassionate therapies to great effect (2, 10). However, despite progress in understanding the ecology and evolution of phage steering *in vitro* (3, 12-14) and *in vivo* (7, 15, 16), we are still in the early phases of a scientific understanding of the phenomenon as a therapeutic tool. There are two main challenges to employing phage steering as a predictable alternative to other therapies. First, using phages to steer pathogenic phenotypes relies on the development of resistance to the phage treatment and that, through a trade-off, resistance somehow compromises the bacterial population. Bacteria can evolve resistance to phage by mutation of receptors on the bacterial cell outer membrane, and, conversely, the evolution of bacterial resistance provides an opportunity for phage mutants able to use modified receptors to exploit otherwise resistant bacteria. Phages and bacteria readily coevolve *in vitro* through reciprocal selection for infection and resistance phenotypes (for review, see (17)), respectively, and there is some evidence that bacterial pathogens can coevolve with phage *in vivo* (e.g., (18)). Nevertheless, it is unclear how coevolution may impact therapeutic outcomes. Specifically, it will be important to know the conditions under which coevolution occurs and whether/how this could affect the evolution of the trade-offs required for phage steering to be effective.

Second, we need to assess how reliable and repeatable phage steering is *in situ*. This is particularly important for phage-antibiotic combinations, as they are likely to become the most frequent form of phage therapy (19). Previous *in vitro* studies have shown that combined phage-antibiotic introductions are more effective than either separately at reducing bacterial densities (20-23). Moreover, in contrast to *in vitro* investigation, we know little about what determines therapeutic outcomes *in situ*, where disease ecosystem structure (1, 24) and immune system dynamics (4, 6) can play important roles. In cases where the phage selects against antibiotic resistance, the expected outcome of combinations will likely depend on the strength of selection pressure and rate of any coevolution between the phage and their bacteria hosts. Recent modelling work (4) suggests that nonlinear synergy between phage, antibiotic, and the innate-immune system may be required for therapeutic success.

In the present study, we investigated the outcomes of phage steering on antibiotic resistance in the pathogenic bacterium *Pseudomonas aeruginosa* (PAO1) both *in vitro* and *in vivo*. Specifically, we tested the sustainability of the steering effect over coevolutionary time scales (e.g., (25)). Briefly, in experimental microcosms, PAO1 bacteria were continuously cultured with phage OMKO1 either in the absence or presence of one of each of three antibiotics. After 10 serial transfers we determined levels of antibiotic resistance in the coevolved bacteria. In a next step, we evaluated the efficacy of both single or combined introductions of phage and the antibiotic erythromycin in an *in vivo* system by using *Galleria mellonella* (wax moth) larvae as hosts for the bacteria. We monitored wax moth survival and determined levels of antibiotic resistance in the surviving bacteria. We found clear evidence of phage impact on bacterial hosts and a phage steering effect of increased bacterial sensitivity for all three antibiotics tested *in vitro*. Importantly, phage reduced antibiotic resistance levels when treated simultaneously with erythromycin *in vivo*. Our results suggest that steering phages are effective either when used as stand-alone therapies and when used in combination with antibiotics.

## Materials and Methods

### Bacterial culture and phage preparation

*Pseudomonas aeruginosa* (Washington PAO1) was grown at 37 °C (constant shaking at 200 rpm) in King’s B (KB) medium (26). For phage amplification, 10 µl of purified OMKO1 phage (3) was added to mid-log growing PAO1, then incubated overnight (18 hours) with intermittent shaking (1 min every 30 mins) at 37 °C (Bülher compact shaker, model KS-15 Control). 10% vol/vol of chloroform was added, the mixture vortexed and centrifuged (1 min 10000 RCF) to separate the phases. The supernatant containing phage was pipetted into a sterile 2 ml Eppendorf and stored at 4 °C. For the complete phage preparation protocol, see (25).

### Experiment 1: Short-term *in vitro* selection for phage resistance

In this experiment, we generated phage-resistant bacteria and tested for a correlated decrease in antibiotic resistance, previously described by (3). PAO1 bacteria from a -80°C freezer stock was grown on KB agar plates overnight at 37 °C. Three single colonies were isolated with a sterile loop, each inoculated individually into 6 ml of KB media, and incubated for 4 hours shaking (200 rpm) at 37 °C. After 4 hours, 10 µL of OMKO1 (*c* 1 × 10^6^) were added, and the three phage-bacteria replicate cultures incubated with intermittent shaking (1 min every 30 mins) at 37 °C for 18 hours. The surviving bacteria were streaked onto a KB agar plate and incubated at 37 °C overnight. Phage resistance was confirmed by cross streaking the bacteria against the phage (27).

To assess the level of antibiotic resistance of the phage-resistant bacteria, the minimum inhibitory concentration (MIC) (28) was determined for three antibiotics with different modes of action (tetracycline, erythromycin, ciprofloxacin). Briefly, phage-resistant bacteria were re-streaked onto KB agar plates overnight, then grown in liquid KB overnight to ensure phage-free cultures. Optical densities of these cultures were measured at a wavelength of 600 nm (OD_600_) and then re-suspended to a uniform OD_600_ of 1.0. In the same way, we prepared ancestral (phage-susceptible) bacterial cultures. In 96-well microtiter plates, a 2-fold dilution series of each antibiotic was prepared, starting at 800 µg/ml and finishing at 6.25 µg/ml for tetracycline and erythromycin, and 10 µg/ml to 0.156 µg/ml for ciprofloxacin. Bacteria were added to each well to a final OD_600_ of 0.05 and incubated at 37 °C for 18 hours. Bacteria were considered as having grown if the OD_600_ was above 0.1 after 18 hours. The MIC was determined as the lowest antibiotic concentration at which no bacterial growth occurred. Three individual colonies were isolated from each replicate culture and three from the ancestor, and their MIC determined for three ‘technical’ replicates. The mean over the technical replicates was used for further analysis.

### Experiment 2: *In vitro* bacterial (co-)evolution

We tested longer-term (10 transfers over 20 days) bacterial evolutionary responses to antibiotics and phage. Bacteria were exposed to a single antibiotic (tetracycline, erythromycin or ciprofloxacin), in the presence of the phage OMKO1, or in the presence of both antibiotic and phage. In a control treatment, neither antibiotic nor phage were added. The experiment was conducted in 2-mL microcosms of KB medium in 24 well microtiter plates, incubated at 37 °C with intermittent 200 rpm orbital agitation for 1 min every 30 mins (Bülher compact shaker, model KS-15 Control). For each of the four treatments, 4 replicate lines were initiated in separate microcosms, for a total of 32 lines across all treatments (4 × control + 4 phage + 3 × 4 Antibiotic-only + 3 × 4 Phage plus antibiotic). Each microcosm was inoculated with 20 μL of PAO1 (*c.* 5 × 10^6^ cells) from an overnight population initiated from a single ancestral *P. aeruginosa* PAO1 colony. For treatments with phage, 5 μL of the ancestral phage stock (*c.* 5 × 10^6^ phage particles) were added together with the inoculated bacteria, producing a 1:1 ratio of bacteria and phage. For treatments with antibiotics, we used half the MIC at the start and again at each serial transfer (200 ug/ul tetracycline, 100 ug/ul erythromycin, 0.5 ug/ul ciprofloxacin). For each microcosm, a 20-μL sample of culture (containing both bacteria and phage) was transferred to a new microcosm with fresh KB medium every 48 hours, for a total of 10 transfers (*c*. 70 bacterial generations for controls). At each transfer, a 50 μL sample was stored in 50% glycerol at -80 °C for subsequent analysis. At the end of the 10 transfers, we determined the antibiotic resistance of the evolved bacteria (MIC, as described above) for 3 arbitrarily chosen colonies from each replicate line. MIC measurements were taken for 3 technical replicates per colony and averaged for statistical analysis.

### Experiment 3: Host mortality and antibiotic resistance evolution in the wax moth infection model

We tested the combined effects of phage and antibiotic on bacteria-induced wax moth mortality and bacterial antibiotic resistance evolution. For the wax moth mortality assay, we used a modified protocol based on (29). Four experimental treatments were established. Late-instar larvae were injected with a total of 20µl containing ancestral PAO1 bacteria (5 × 10^2^ CFU), supplemented with phage only (5 × 10^3^ PFU), the antibiotic erythromycin only (concentration: 100 µg/ml suspended in PBS), both phage and antibiotic, or neither phage nor antibiotic (= positive control). This fully factorial experiment was replicated four times, within each replicate run we used 6 larvae per treatment, giving a total of 96 larva. In an additional negative control treatment, 18 larvae were injected with 20 µl phosphate saline buffer (PBS), without bacteria. Larvae were kept at 37 °C for the two days following injection and the survival of the treated larvae was checked at 6-h intervals for the first 24h, then at 12-h intervals. Larvae were considered dead when 3 successive stimulations along their ventral surface produced no response. All dead larvae were stored intact in 25% glycerol at -80 °C at the end of the experiment at 48 hours.

Bacterial antibiotic resistance was determined from 4 wax moth larvae in each of two arbitrarily selected replicate lines for each treatment (therefore, 8 wax moths analysed per treatment). Each individual larva was homogenized at 48 hours by pestle sticks (Bel-Art instruments). The resulting homogenates were then serially diluted and plated onto *Pseudomonas* Isolation Agar (PIA, DifCO). The MIC was determined for three arbitrarily selected single colonies per wax moth analysed, and the mean recorded as described for Experiment 2.

## Statistical analysis

For the initial screen of phage-resistant PAO1 (Experiment 1), MIC data of bacterial resistance to the three antibiotics were analysed by a t-test (Welch two-sample test), comparing evolved phage-resistant isolates with phage-susceptible ancestral bacteria. Analysis of Variance (R 3.01 base package, supplemented with the ‘car’ and ‘lsmeans’ packages using the Tukey adjustment) was employed for the MIC data from Experiment 2 (*in vitro* evolution). A fully factorial model of phage and antibiotic treatments was fitted, with replicate selection line nested in the interaction term.

To analyse variation in larval mortality associated with bacteria in Experiment 3, we fitted a parametric survival model with time to death as the response variable, using the Weibull distribution function (R packages ‘survival’ and ‘survminer’). A Weibull distribution function was chosen for having the lowest AIC score in comparison to models with other functions (e.g., lognormal, exponential). Right censoring was used for cases of larvae surviving the full 48h post-injection. Phage and antibiotic treatments (and their interaction) were fitted as explanatory factors in a factorial model, as above. The identity of experimental replicate run was removed from an initial full model, since it explained a non-significant portion of the variance (p > 0.1). Its removal did not bias our results for the remaining effects in the model. We further analysed variation in MIC among bacteria isolated from dead larvae, with phage and antibiotic treatments (and their interaction) as explanatory factors. MIC data in all analyses were square root transformed to meet model assumptions. These analyses were carried out in JMP 14 (SAS).

## Results

### Experiment 1: Short-term selection for phage resistance leads to reduced MIC *in vitro*

As shown previously (3, 11), PAO1 bacteria that had become resistant to phage OMKO1 after 1 day of exposure were found to have significantly lower antibiotic resistance (Fig. 1). For all three antibiotics tested, the group of phage-resistant bacteria isolates had greater than 70% lower MICs than the phage-susceptible ancestral bacteria (ciprofloxacin: t_2.04_ = 14.9, *p* = 0.004; erythromycin: t_3.2_ = 8.5, *p* = 0.013; tetracycline: t_3.92_ = 16.63, *p* = 0.004).

**Figure 1.**
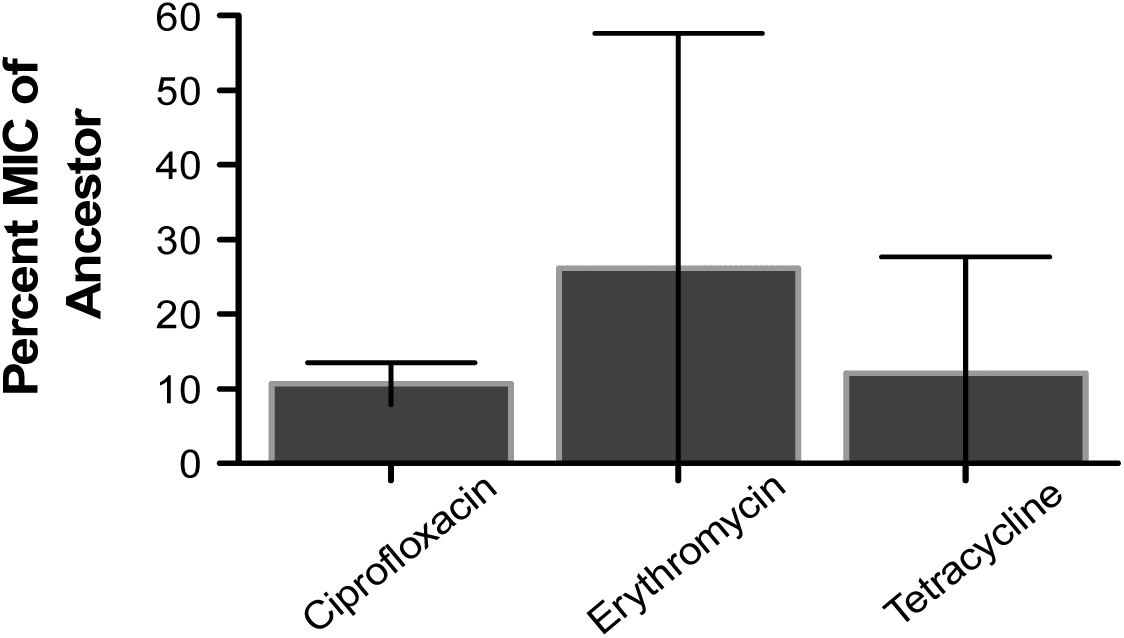
Percent (± 95% CI) of MIC of phage-resistant bacteria tested for three antibiotics relative to ancestral phage-susceptible bacteria. PAO1 bacteria resistant to OMKO1 were obtained after 24 h of *in vitro* growth in the presence of the phage. MIC levels were determined for each of three antibiotics; all three were significantly reduced from the initial ancestor level (t-test, Welch two sample test).

### Experiment 2: Long-term exposure to phage prevents the emergence of antibiotic resistance

Analyses of bacterial resistance (MIC) after 10 transfers revealed a general negative effect of phage treatment on MIC (significant main effects; Table 1). Bacteria that had been co-cultured with phage showed greater than 60% reductions in resistance to all three antibiotics compared to bacteria that had not been exposed to phage (Fig. 2). In contrast, there was no clear general signal of antibiotic treatments (Table 1; Fig. 2). Erythromycin significantly increased resistance (main effect of antibiotic treatment), tetracycline had no significant effect, while the effect of ciprofloxacin depended on the presence of phage (significant antibiotic x phage interaction; Table 1).

**Table 1.** Analyses of variance of antibiotic resistance (minimum inhibitory concentration, MIC) for bacteria isolated from selection lines undergoing treatments with and without exposure to phage OMKO1 or each of three antibiotics (erythromycin, tetracycline, ciprofloxacin). All treatment effects were tested against the selection line effect. Significant effects in bold.

| Source | d.f. | Erythromycin |  |  | Tetracycline |  |  | Ciprofloxacin |  |  |
| --- | --- | --- | --- | --- | --- | --- | --- | --- | --- | --- |
|  |  | MS | F | *p* | MS | F | *p* | MS | F | *p* |
| Phage | 1 | **679.2** | **42.38** | **<0.0001** | **277.2** | **66.8** | **<0.0001** | **9.0** | **164.2** | **<0.0001** |
| AB | 1 | **1315.4** | **82.07** | **<0.0001** | 9.37 | 2.26 | 0.1586 | 0.12 | 2.34 | 0.15 |
| Phage x AB | 1 | 4.1 | 0.25 | 0.62 | 0.03 | 0.007 | 0.9329 | **0.83** | **15.23** | **0.002** |
| Selection line (phage, AB.) | 32 | 192.3 | 0.72 | 0.71 | 4.14 | 0.62 | 0.8077 | 0.05 | 0.87 | 0.5810 |
| Residual | 54 |  |  |  |  |  |  |  |  |  |

**Figure 2.**
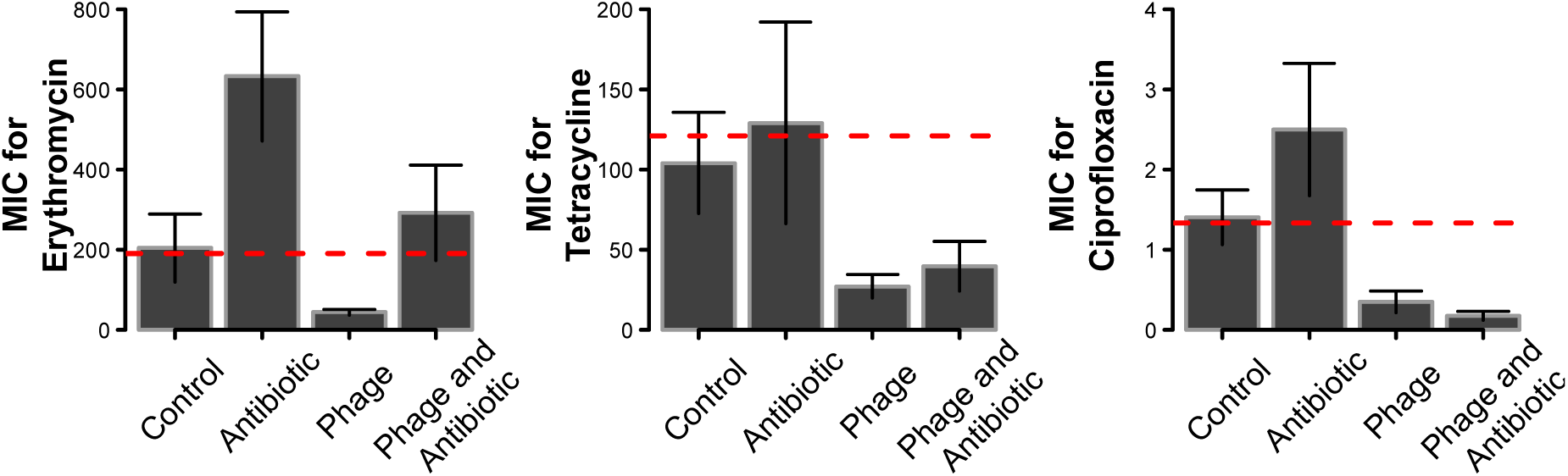
Levels of antibiotic resistance (minimum inhibitory concentration, MIC ± 95% CI) of bacteria evolved in the presence of an antibiotic only, phage only, or both phage and antibiotic combined. a) Erythromycin. b) Tetracycline, c) Ciprofloxacin. Bacteria were isolated after 10 serial transfers (20 days) of treatment and their MICs determined. Bacteria from antibiotic treatments were only tested against the same antibiotic; bacteria from phage-only treatments were tested against all three antibiotics. Phage OMKO1 was able to either reduce the emergence of resistance or sensitize the bacteria to the antibiotic. Dashed red line is the mean MIC for the ancestor.

Detailed pairwise treatment comparisons of MIC values showed two patterns of phage action. First, adding phage together with the antibiotic reduced bacterial resistance relative to antibiotic-alone treatments (all *p* < 0.002; Fig. 2a-c). In at least one case (ciprofloxacin), phage and antibiotic clearly had non-additive effects (significant phage x antibiotic interaction, Table 1), such that the MIC in the combined treatment was more similar to that of the phage-alone treatment (rather than the average of the two single treatments). Second, all three phage-alone treatments and two combined phage-antibiotic treatments reduced MIC levels below that of bacteria from untreated controls (all *p* < 0.05; Fig. 2). Only for erythromycin did we observe some level of antibiotic resistance evolution in the combined treatment, but this was not significant (*p* > 0.05) (Fig 2a). Taken together, these patterns indicate that phages not only prevented antibiotic resistance evolution, but even increased susceptibility of the bacteria to ciprofloxacin and tetracycline (Fig. 2b,c).

### Experiment 3 (survival assay): Phage reduces bacteria-related larval mortality

Bacterial injection reduced larval survival relative to the PBS-injected negative control group (1 dead larva out of 18, data not shown) (Figure 3). In the absence of phage or antibiotic, bacterial infection was near 100% lethal within 24h (a single larva survived to 48h with erythromycin). Even with anti-bacterial treatments (phage and/or antibiotic) more than 40% of the larvae died within 48h.

**Figure 3.**
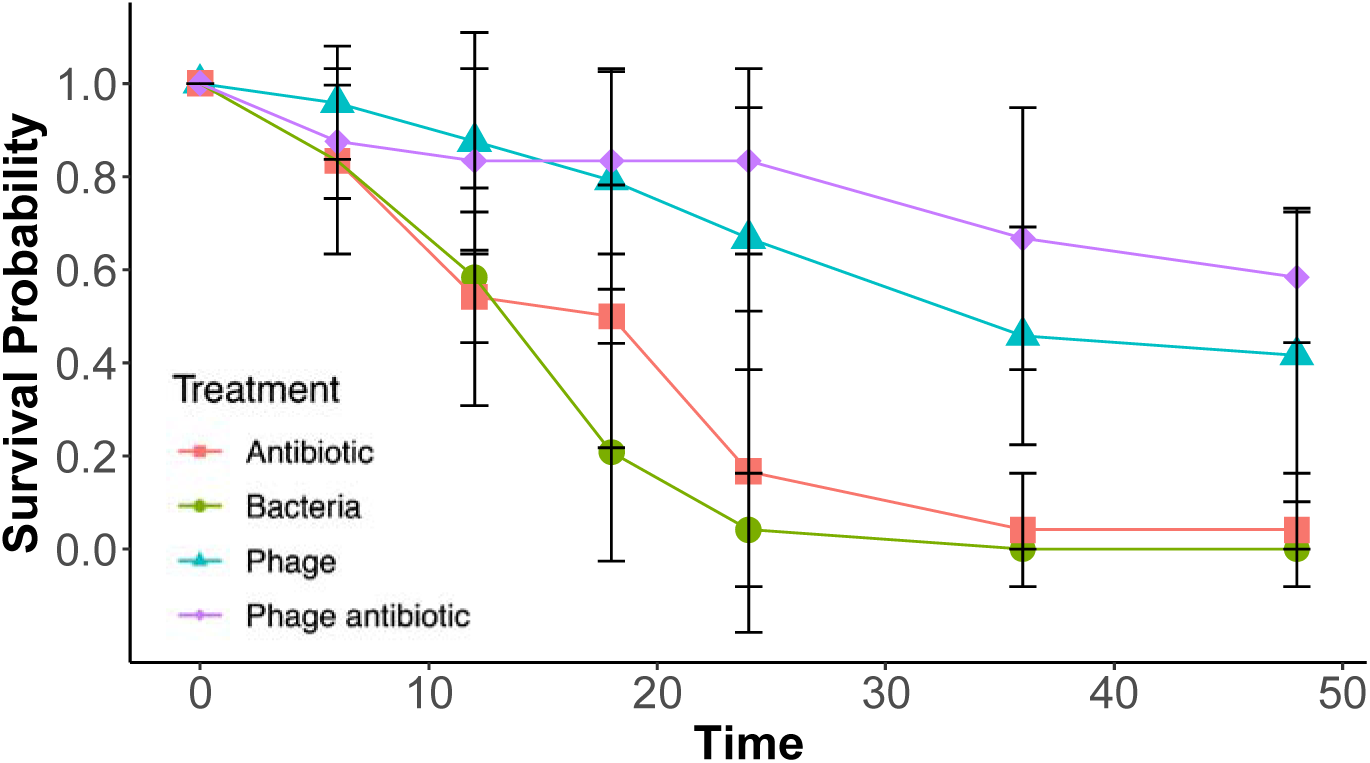
Mean (± 95% CI) proportion of surviving wax moth larvae under the 4 different treatments. Larvae were infected with a lethal dose of PAO1 and observed for 48 hours. All 3 treatment lines provided a greater mean survival time. Both Phage alone (Blue Triangle) and Phage plus Erythromycin (Purple diamonds) produced an additive survival rate when compared to Erythromycin alone (Red Squares). Mean survival of 24 replicates error bars are 95% CI.

The addition of the phage OMKO1 and of the antibiotic each significantly increased larval survival (phage: Wald χ^2^ = 58.8, *p* < 0.0001; antibiotic: χ^2^ = 5.1, *p* = 0.0242; Figure 3). However, while the phage treatment prolonged survival by *c.* 20h and left more than 50% of the larvae alive at the end of the 48-hour assay, the positive effect of the antibiotic was only marginal (4h longer life; 96% mortality by 48h) and its significance depended on the distribution function chosen (e.g., p > 0.2 for a lognormal survival function). There was no significant interaction between antibiotic and phage treatment (χ^2^ < 1, *p* > 0.8), indicating that the two anti-microbial agents acted additively. Thus, the combined injection of phage and antibiotic was most successful at prolonging larval survival, even though it was not significantly different from the phage-alone treatment (pairwise contrast p > 0.2).

### Experiment 3 (MIC assay): Phage maintains selection against antibiotic resistance *in vivo*

Factorial ANOVA showed a highly significant overall effect of phage treatment on the MIC of evolved bacteria (F_1,28_ = 132.7, *p* < 0.0001), whereas the main effect of the antibiotic treatment and the antibiotic x phage interaction were both non-significant (F_1,28_ = 0.4, *p* > 0.5, and F_1,28_ = 0.5, *p* > 0.4, respectively). While the antibiotic alone led to only a slight increase in MIC, the action of the phage strongly reduced the MIC of the surviving bacteria (Figure 4). Thus, like in the *in vitro* experiment, the presence of phage increased susceptibility of the bacteria to the antibiotic, resulting in lower MICs than in the untreated control lines.

## Discussion

We present evidence of a phage associated reduction in antibiotic resistance in bacterial populations of *Pseudomonas aeruginosa*, both *in vitro* and *in vivo*. In line with previous work (3), phage OMKO1 reduced antibiotic resistance in the majority of treatment conditions (by up to 90% Fig.1) and impeded the emergence of resistance in those it could not reduce (Fig 2). *In vivo* assays largely confirmed the *in vitro* experiments, indicating that OMKO1 both reduces antibiotic resistance and when combined with certain antibiotics can have a larger impact on target bacteria compared to either antimicrobial used separately, thereby improving larval host survival.

**Figure 4.**
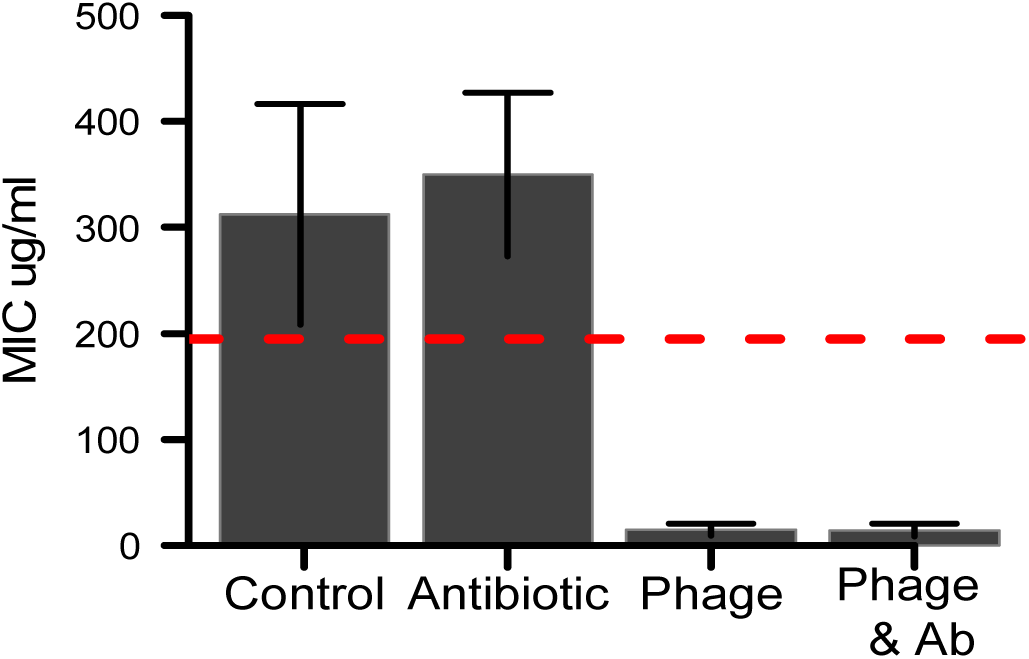
Mean MIC (± 95% CI) of recovered PAO1 bacteria from wax moth infection assay. Bacteria were recovered from 8 wax moth larvae per treatment. MIC was determined using serial dilution. Both the Phage and Phage + Antibiotic (erythromycin) line produced clear reductions in the MIC level post infection.

The PAO1 isolate used in this study was determined to be clinically resistant to ciprofloxacin by both the CLSI and BSAC breakpoints (30)(31). Neither erythromycin nor tetracycline are routinely used clinically for *Pseudomonas aeruginosa* treatment and therefore do not have established clinical resistance levels (CLSI and BSCA do not set breakpoints). Nevertheless, the ancestor MICs for both erythromycin and tetracycline reported here were in line with previous studies (32, 33), and post phage treatment MICs for both were significantly reduced (Fig 1). All three antibiotics are known to effluxed by OprM-MexXY systems (32, 34, 35), which appear to be targeted by phage OMKO1.

The long-term (20-day) experiment yielded a number of interesting and promising results. OMKO1 was able to significantly reduce resistance or impede the emergence of resistance in all three antibiotic treatments even in the presence of the antibiotic (Fig 2). This demonstrates that OMKO1 might be used as an adjuvant in antibiotic treatments by selecting for antibiotic sensitivity and contributing along with the immune system to bacterial clearance (4). Such treatments have been undertaken as expanded access compassionate care cases and are currently being analysed.

As a proof of principle for such treatments and to test the interaction of phage, antibiotic and innate-immune system we performed a simple infection assay using wax moth larvae. The larvae were injected with a lethal quantity of bacteria with or without erythromycin and/or phage, and then observed for 48 hours. The mean survival time increased for all three treatment conditions compared to bacteria injected alone (Fig 3). Fitting a parametric survival model however failed to show a synergistic effect of phage and antibiotic, and rather gave evidence for an additive response. Our results nevertheless support the basic findings of other studies showing improved treatment outcomes with phage for a range of *in vivo* systems ranging from *C elegans (15)* to mammals including humans (7, 11, 36-39). Our work is the first to our knowledge to show a reduction in resistance in any *in vivo* system. This result taken together with the *in vitro* experiments suggests that phage steering can occur *in vivo* and could be evolutionary stable during the course of treatment.

Although we have shown that OMKO1 can reduce bacterial numbers and maintain antibiotic sensitivity even in the presence of the latter, several unanswered questions remain. Of particular interest is the extent to which the phage maintains selection on the target, in this case the *oprM*-*mexXY* system. This could be investigated by examining efflux pump genetics during a similar long-term experiment, where both phage and bacteria potentially (co)evolve and are independently allowed to evolve against the antibiotic (40). Coevolution under different environmental pressures may shift dynamics between ‘arms race’ and ‘fluctuating’ with implications for phage resistance (41, 42). Introducing other bacterial species that capture the multispecies nature of many infections may also change how resistance develops (43), shifting selection from the receptor target to other resistance mechanisms such as CRISPR-cas (43). Theoretical work however indicates that selection will often result in receptor modification (44).

The use of wax moths as *in vivo* experimental systems has two main drawbacks. First, the immune system is limited to an innate response. Although an innate response can be key to successful phage therapy (4, 6), the influence of the adaptive response (as would be the case in human subjects and other animal models such as mice) is currently unknown. Second, the infection and its treatment were introduced simultaneously, which (possibly excepting prophylaxis) is clearly irrelevant to real treatments. This simultaneous administration likely means that the bacteria had little time to grow and adapt to the infection or to form structures such as biofilms that may limit the ability of phage to infect (45). Physical limitations of the wax moth model mean that it is difficult establish an infection and then apply the treatment, as death can be rapid and multiple injections are likely to produce non-infection-related mortality. Future study should investigate different delays between infection and the introduction of antibiotics and phages in murine models, where immune responses are more similar to human *in situ* contexts.

Our study examined whether phage OMKO1 could steer antibiotic resistance successfully during *in vitro* experimental evolution and in an animal model, without regard to the spectrum of phage-resistance and drug-resistance mutations underlying phenotypic changes in bacterial populations. These experiments considered the underlying genetics as unopened ‘black boxes’, analogous to treatment situations such as chemotherapy administered against cancer and antibiotic therapy targeting bacterial infections, where the mutational responses to therapy are typically neither elucidated nor monitored. These established therapies assume the mutational spectra for drug resistance can vary but should generally not impede a successful treatment outcome in the patient. Measuring at the site of an infection is impractical, thus, these therapies could be considered gambles, which presume the beneficial phenotypic responses will win out over unfavourable clinical outcomes based on *in vitro* or *in vivo* experimental evidence. Similarly, the widespread use of phage therapy may move forward by relying on confidence that the treatment can be delivered safely and effectively, while simultaneously ignoring many of the underlying mechanistic details. Our ongoing work examines the generalities for *P. aeruginosa* resistance to phage OMKO1, by mapping the spectra of fitness and phenotypic effects for these mutations in PAO1 and for strains isolated in the clinic.

In summary, as expected OMKO1 was able to achieve a reduction in MIC for all three antibiotics (Fig 1). Our results support the action of a trade-off between the reduction of MIC and phage resistance (Fig 2), suggesting that the phage was able to maintain selection against resistance over long term, co-evolutionary time scales even in the presence of antibiotics. This effect was confirmed *in vivo*, where the phage improved survival and reduced antibiotic resistance (Figs 3, 4).

## Acknowledgments

MEH thanks the McDonnell Foundation for funding (Studying Complex Systems research award 220020294).

